# Chromatin structure changes in *Daphnia* populations upon exposure to environmental cues – or – The discovery of Wolterecks “Matrix”

**DOI:** 10.1101/824789

**Authors:** Ronaldo de Carvalho Augusto, Aki Minoda, Oliver Rey, Céline Cosseau, Cristian Chaparro, Jérémie Vidal-Dupiol, Jean-François Allienne, David Duval, Silvain Pinaud, Sina Tönges, Ranja Andriantsoa, Emilien Luquet, Fabien Aubret, Mamadou Dia Sow, Patrice David, Vicki Thomson, Déborah Federico, Dominique Joly, Mariana Gomes Lima, Etienne Danchin, Christoph Grunau

## Abstract

Phenotypic plasticity is an important feature of biological systems that is likely to play a major role in the future adaptation of organisms to the ongoing global changes. It may allow an organism to produce alternative phenotypes in responses to environmental cues. Modifications in the phenotype can be reversible but are sometimes enduring and can even span over generations. The notion of phenotypic plasticity was conceptualized in the early 20^th^ century by Richard Woltereck. He introduced the idea that the combined relations of a phenotypic character and all environmental gradients that influence on it can be defined as “norm of reaction”. Norms of reaction are specific to species and to lineages within species, and they are heritable. He postulated that reaction norms can progressively be shifted over generations depending on the environmental conditions. One of his biological models was the water-flee *daphnia*. Woltereck proposed that enduring phenotypic modifications and gene mutations could have similar adaptive effects, and he postulated that their molecular bases would be different. Mutations occurred in genes, while enduring modifications were based on something he called the *Matrix*. He suggested that this matrix (i) was associated with the chromosomes, (ii) that it was heritable, (iii) it changed during development of the organisms, and (iv) that changes of the matrix could be simple chemical substitutions of an unknown, but probably polymeric molecule. We reasoned that the chromatin has all postulated features of this matrix and revisited Woltereck’s classical experiments with *daphnia*. We developed a robust and rapid ATAC-seq technique that allows for analyzing chromatin of individual daphnia and show here (i) that this technique can be used with minimal expertise in molecular biology, and (ii) we used it to identify open chromatin structure in daphnia exposed to different environmental cues. Our result indicates that chromatin structure changes consistently in daphnia upon this exposure confirming Woltereck’s classical postulate.

## Introduction

We are exploring here a classical battlefield of evolutionary biology. In a now landmark presentation in 1909, at the annual meeting of the German zoological society in Frankfurt, and at a time when oral presentations were still the major avenue for scientific exchange, “*Mr. R. Woltereck (Leipzig)*” exposed his ideas on *Artveränderungen* (change of species). The very same year, the scientific community had been captured by the publication of the German version of university lecture materials by the Danish botanist W. Johannsen, going back to 1903 (Johannsen 1909). Johannsen had introduced the terms of *phenotype* and *genotype* to separate the outer impressions we have of an organism from the heritable components it has inside. He also had introduced the notion of “pure lines” on which selection would be powerless since offspring of selected phenotypes would still produce the same range of phenotypes. His work was based on the mutation theory of de Vries (Vries & MacDougal 1905) who had stated that the characters of organisms are made of distinct units that change spontaneously, salutatory, and relatively rarely. de Vries had called these changes *Mutations*. Mutations were in his eyes heritable and could be selected for. Woltereck’s criticism was that the environment had nno influence on de Vries “mutations” or the “exact science of heritability” of Johannsen. This was counterintuitive to Woltereck and many fellow scientists who saw that the environment definitely had an impact on the phenotype. His battle horse became *daphnia*, easy to handle and cheap to maintain (contemporaries will understand the attractiveness of the system). Woltereck reasoned that, in response to Johannsen, further “analytical” experiments should also be done with “pure lines” (clonal lineages in modern terms), and with quantitative characters to investigate the role of the “milieu” (environment) on the character (Woltereck 1909). Woltereck recorded morphological measures, in particular the relative head length of his *daphnia* lines, depending on environmental conditions such as temperature and nutrition. These early studies paved the way to a subsequent rich literature that has documented the amazing property of *daphnia* to modify their phenotypes at the morphological, physiological, behavioral and more recently at the molecular levels in response to a large panel of environmental stressors including diet, pollution, heavy metals, and predator kairomones (reviewed in (Riessen 2011, Harris *et al.* 2012)). He called these relations of the phenotype on an environmental gradient *Phänotypenkurve*. The combined relations of a phenotypic character and all environmental gradients that influences it, he defined as *Reaktionsnorm* or “norm of reaction”. According to him, norms of reaction are specific to species and to lineages within species, they are heritable and based on (in his opinion are equal to) the genotype. He postulated that reaction norms can progressively be shifted over generations depending on the culture conditions of *daphnia* (Woltereck 1909). Later, in his 1932 textbook (Woltereck was a lecturer at the University of Leipzig), he extended this view to the notion that species should be defined by identical norms of reaction (Woltereck 1932). This is a remarkable concept in the light of current difficulties to define species boundaries by phenotypic similarity, reproductive isolation or DNA sequence similarity. He also expanded the concept of the norm of reaction to three types of norms: 1st order (*Modifikationen*), 2nd order (*Kombinationen*), and 3rd order (*Dauerinduktion* and gene mutations). *Modifikation* was a textbook term in the 1920-30s and is equivalent to phenotypic variation. We will focus here on the 3rd order norms of reaction. Woltereck borrowed the term *Dauerinduktion* from Victor Jollos who had coined in the early 1900s; the term *Dauermodifikation* or “enduring modifications” (Jollos 1939), to describe phenotypic changes that could be provoked by environmental stimuli, would persist for a few generations and then revert.

Interestingly, Woltereck considered enduring modifications and gene mutations somehow similar. Nevertheless, he proposed that the molecular basis would be different. Mutations occurred in genes, while enduring modifications were based on something he called the *Matrix*. He suggested that this matrix was associated with the chromosomes (“… *chromosomes are matrix plus gene…*”), that it was heritable, changed during development of the organisms, and that changes of the matrix could be simple chemical substitutions of an unknown, but probably polymeric molecule. (More on Woltereck’s work at https://embryo.asu.edu/pages/richard-wolterecks-concept-reaktionsnorm and (Nicoglou 2017)). The phenomenon that organisms change their appearance as a function of environmental cues and/or during development is today rather called phenotypic plasticity, a term introduced in the 1960s. Mayr (Mayr 1963) used “polyphenism” to distinguish environmentally induced phenotypic variation from those that he believed were genetically determined (polymorphisms). Two years later, Bradshaw termed the amount by which the expression of an individual genotype can be modified by its environment as “plasticity” (Bradshaw 1965) and discussed the importance of plasticity for evolution. Nowadays, the importance of developmental and environmental plasticity for the generation of phenotypic novelty is still a matter of lively scientific discussion (Levis & Pfennig 2019). But it is increasingly recognized that enduring phenotypic plasticity requires memory effects that can be related to epigenetic mechanisms. The definition of what is epigenetic depends very much on the scientific context in which the term is used (Nicoglou & Merlin 2017). Here we will use it for any chromatin modification affecting gene expression, whether it is heritable or not (Nicoglou & Merlin 2017) and we will show that it is related to Woltereck’s matrix.

In the last two decades, several studies have started to identify the molecular basis of such a matrix including non-coding RNAs, covalent modifications at the histone tails and DNA methylation. All of these mechanisms together constitute the epigenetic information that allows the remodeling (and maintenance) of chromatin structure and ultimately of phenotypes under environmental influence. In this regard, the global level of DNA methylation of *Daphnia magna* was found to be largely affected after exposure to abiotic (e.g. Zinc; (Vandegehuchte *et al.* 2010)) and biotic (toxic cyanobacterium *Microcystis aeruginosa*; (Asselman *et al.* 2017)) environmental toxicants, or to irradiation (Trijau *et al.* 2018). However, no study has yet investigated the effect of environmental stimuli on the genome-wide chromatin structure. Here we argue that what Woltereck called the matrix is nowadays chromatin structure; the bearer of the overall epigenetic information including all epigenetic marks and their complex interactions. Here, we adapted an ATAC-Seq (Assay for Transposase Accessible Chromatin with high-throughput sequencing) (Buenrostro *et al.* 2013) protocol to characterise the overall genome-wide chromatin structure of *Daphnia pulex* in the context of the iconic complex defense response to predation. ATAC-seq works similarly as DNase-seq (DNase I hypersensitive sites with high-throughput sequencing) (Song & Crawford 2010), and determines which genomic regions are accessible to Tn5 transposase (*i.e.* open chromatin regions), especially the regulatory regions. Tn5 transposase inserts Illumina adapter sequences upon accessing the chromatin, which removes the need for additional steps to make the sequencing libraries later. This simple and efficient protocol reduces the enables starting material required, compared to DNase-seq. It also avoids many other steps such as the interaction with antibodies (e.g. ChIP-seq) or chemical treatment (e.g. FAIRE-seq, WGBS) that might introduce bias.

Our results show: (i) that ATAC-seq can be used to characterize chromatin structures of individuals even those that are small and thus with few biological material, making it possible to determine epigenetic polymorphisms relatively easily and at reasonable cost in full populations; and (ii) we deliver evidence that chromatin structure changes upon stimuli from the environment (figure 1).

**Figure 1:**
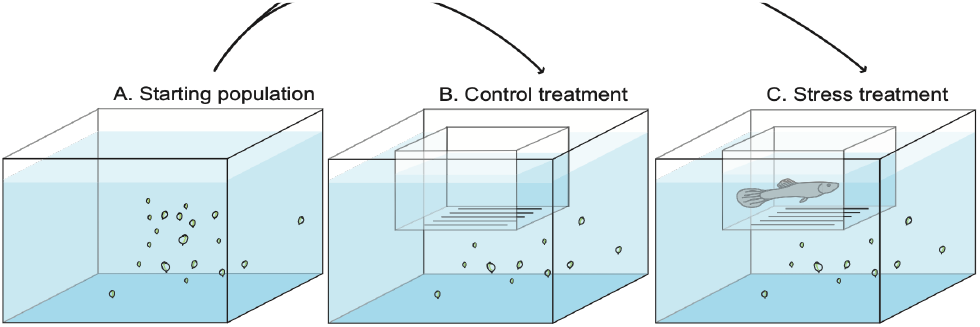
Experimental design used in this study. *Daphnia* were put into a water tank and allowed to acclimate (start population). Then, two experimental tanks were set up following strictly the same design. The only difference was the presence of a predator (a guppy trained to eat daphnia) in the floating plastic fish breeding isolation box in the stress treatment.

This study therefore describes the classical experimental system postulated by Richard Woltereck 100 years ago: the adaptive morphological phenotypic plasticity of *daphnia*.

## Results

### ATAC-Seq can be used on individual Daphnia

Our ATAC-seq procedure delivered reproducible chromatin profiles for individual *daphnia*. Projection of ATAC-seq reads on a metagene profile indicated that Tn5 accessible and thus presumably open chromatin structure occurs at the TSS and in gene bodies (Figure 2, suppl. figure 1).

**Figure 2:**
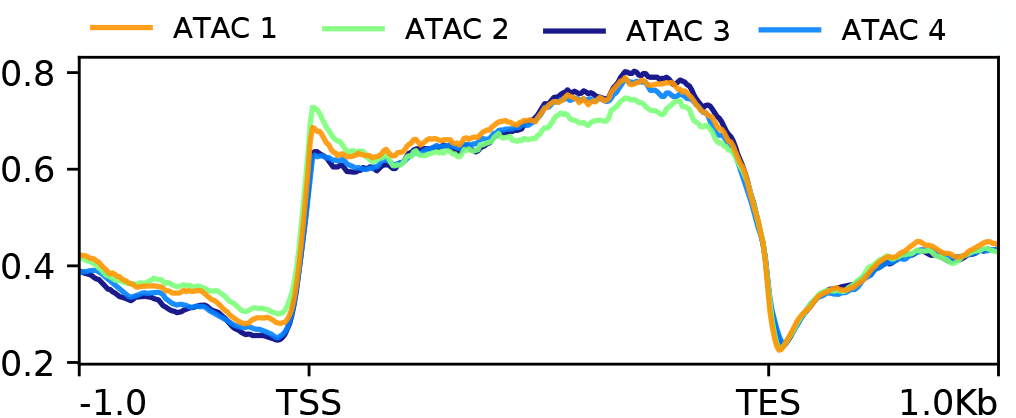
Superposed metagene ATAC profiles of four individual *daphnia* of the start populations. X-axis in base-pairs. TSS = Transcription start site, TES = transcription end site. Y-axis average enrichment of ATAC-seq reads over genes and upstream and downstream regions. Enrichment of accessible chromatin occurs along the entire length of the genes.

We started by comparing populations sampled at the beginning of the experiment (start) to the unexposed (control) population at the end of the exposure time of the experiment. Clustering algorithms built into DESeq2 were used to produce a graphical representation of sample-to-sample distances based on the similarity of their ATAC-Seq patterns (Figure 3). These data indicate that there are 2,362 local differences (3.6% of all 66,194 identified ATAC enrichment regions) at an FDR of 0.05 between the start and the control population (p-value adjusted for multiple testing with the Benjamini-Hochberg procedure).

**Figure 3:**
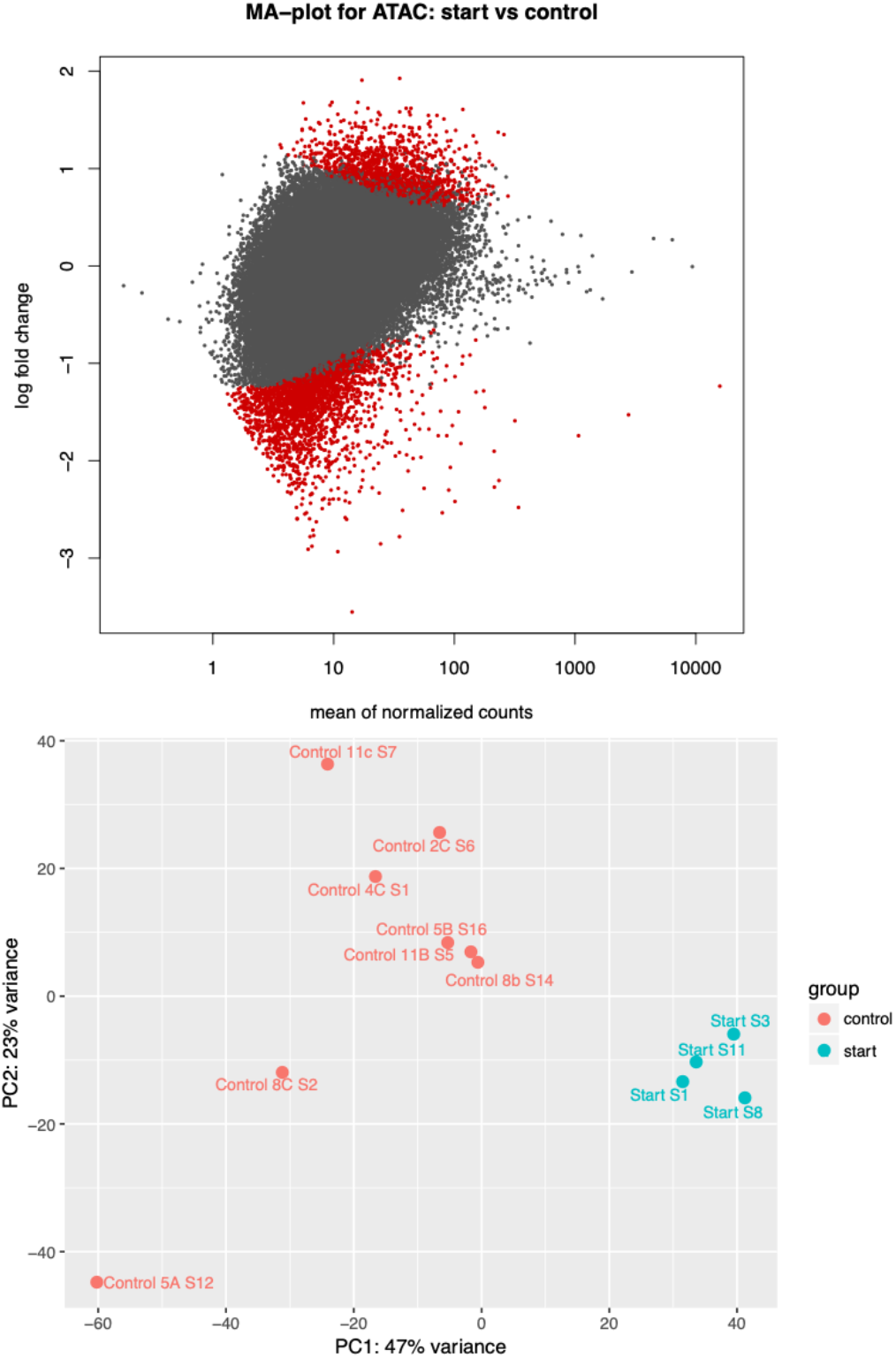
MA-Plot (top) and Principal component analysis of individual *Daphnia* based on their ATAC-Seq profiles (bottom). On the PCA every point represents an individual *daphnia*. Populations are color coded. Samples from the start (blue) and control population (red) cluster clearly.

This suggest that within 20 days (2-5 generations) there was either (i) epigenetic drift from ‘start’ to ‘control’ or (ii) epimutations were induced and/or selected by changes in the water tank environment from ‘start’ to ‘control’.

### Exposure to predator cues leads to morphological differences in Daphnia

Our results show that on average, the (LL-SL)/SL ratio calculated for *daphnia* from the stress treatment (N = 14; Mean = 0.24 ± 0.072) was significantly higher than that of *daphnia* from the control treatment (N = 12; Mean = 0.15 ± 0.039; Mann-Whitney U Test, *U* = 19, Z = −3.32, P < 0.001) (Figure 4). This result confirms the expected induction of anti-predatory morphs in the stress treatment. It is noteworthy that the quantified morphological response to predation pressure observed in the stress treatment most likely reflects a more general response of stressed *daphnia* including morphological, physiological and behavioural changes (Boersma *et al.* 1999) Our first intention in comparing *daphnia* from the two experimental treatments was to confirm that we effectively induced a global response in stressed individuals, these responses having been otherwise much better documented previously (Riessen 1999)

**Figure 4:**
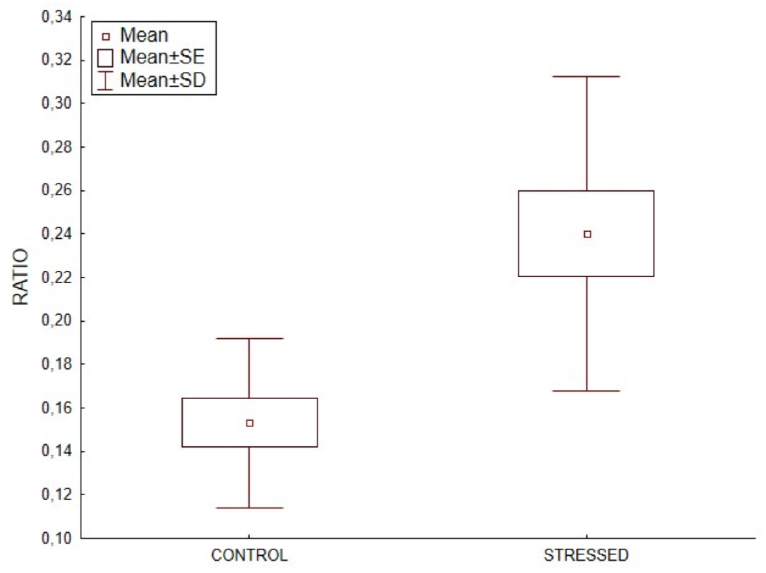
Boxplot of morphometric ratios of (LL-SL)/SL in control and stressed *daphnia* populations. (control: N = 12; stress: N= 14).

### Exposure to predator cues leads to differences in chromatin structure between exposed (stressed) and unexposed (control) Daphnia

Using the DESeq2 procedure described above for ‘start’ vs. ‘control’ we identified 66,194 differences between ‘control’ and ‘stressed’. This is by far too many, and indeed, shifts in MA plots (not shown) indicated that the assumption that is underlying the algorythm used in DESeq2 and the requires that most sites do not change, was violated. Metagene profiles, using the same number of aligned reads over the entire genome, lend further support to the finding that ‘stressed’ samples had on average fewer reads over genes than ‘control’ samples indicating major changes in chromatin structure (Figure 5).

**Figure 5:**
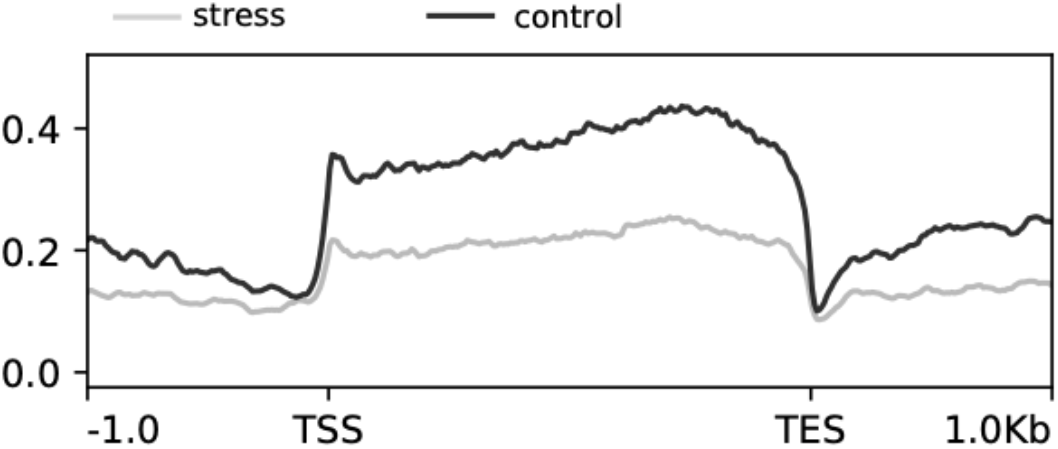
Combined metagene ATAC profiles of stressed and control *daphnia* populations. X-axis in base-pairs. TSS = Transcription start site, TES = transcription end site. Y-axis average enrichment of ATAC-seq reads over genes and upstream and downstream regions.

This also means that there is a large number of regions for which no reads could be recovered in the stressed samples. This is not due to a general lower accessibility of Tn5 to the cells and nuclei because of a thicker cuticle or a similar phenotypic trait because the insert size distribution of start, control and stressed populations are similar (Supplementary file 2). If DNA was more inaccessible in the stressed population we would expect longer fragments. To cope with the general decrease of ATAC-Seq reads in the stressed population, we resorted to ChromstaR, a HMM based software that was developed for ChIP-Seq analysis but that in principle can also be used for ATAC-Seq and is probably less sensitive to zero values. Under the constraints of numerous instances of an absence of data, ChromstaR identified 87 regions that are different between start and control, and stress. All were visually inspected using MACS2 average profiles, normalised by the same number of aligned reads over the genome. Among these 87 regions, ATAC signal was down in stressed samples compared to ‘control and start’ in 45 regions (52%), down in ‘stress and control’ compared to ‘start’ in 16 (18%), up in ‘stress and control’ in 3 (3.4%), and down in ‘control’ in only 1 (1.1%). Seven regions showed a heterogenous pattern on ATAC signals. In 15 regions differences were considered too weak (17%) suggesting that fine tuning of ChromstaR parameters might be necessary (Supplementary file 2). These results are in line with a general decrease in ATAC signal in the stressed samples, i.e. chromatin becomes less accessible and/or less heterogenous. It is interesting to note that for 20 regions adjacent ATAC signals (less than 2kb apart) were detected, lending further support to the idea that chromatin structure changes occur in a controlled fashion.

Clustering of the samples clearly regroups control and stressed samples (Figure 6).

**Figure 6:**
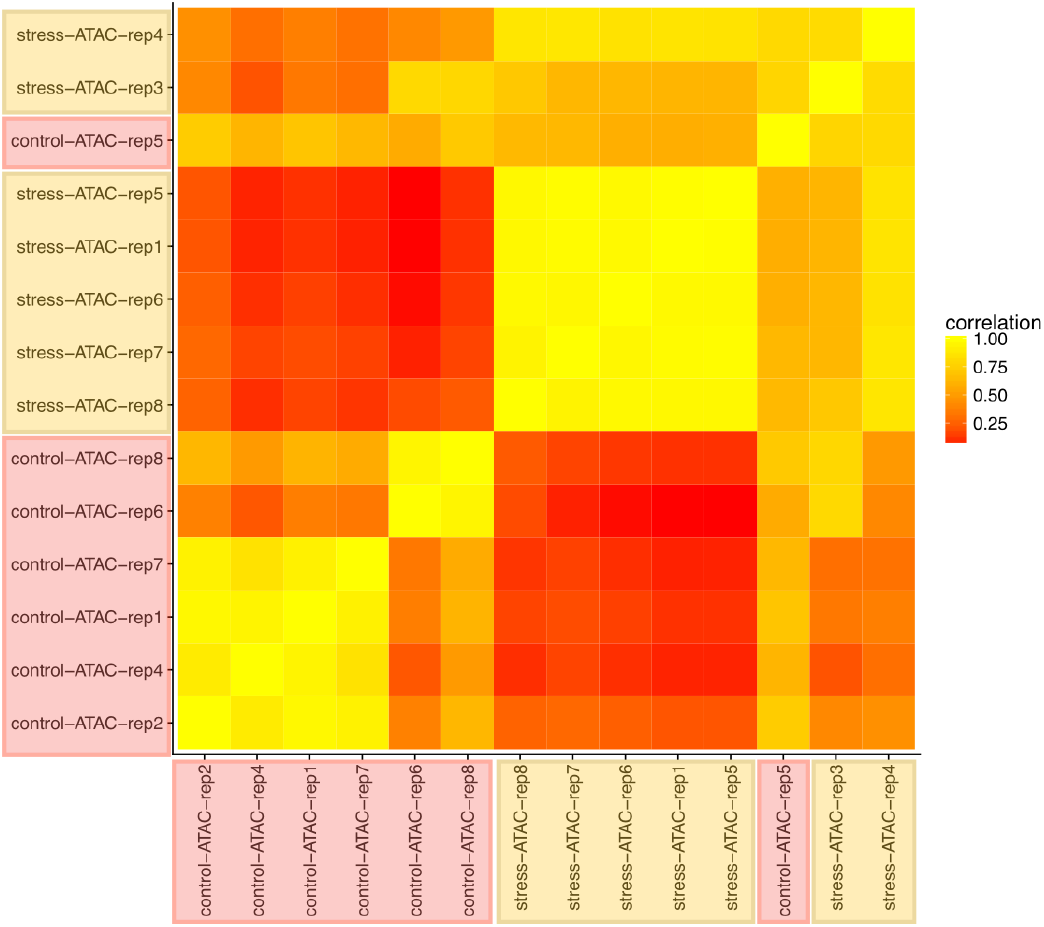
Clustering of individual *daphnia* based on their ATAC-Seq profiles. Heat map indicating similarity in the HMM ChromstaR results. Generally samples from the stressed and the control populations each cluster together.

Another way to cope with presence of many zeroes that could produce a difference between two groups simply because zeroes in one group turn out to be very small values in the other, is to use log transformation; setting an arbitrary low threshold level of accessibility that we do not consider very different from zero. Here we transformed the data with log10(*t*+*x*), where *t* is the threshold of 0.1 and x is the ATAC-seq read count. Doing so we see again that ‘stressed’ are very different from ‘controls’: the distribution has many very small values (including many true zeroes = log10(t+x) = −1) and ‘stress’ mode is slightly shifted to the left. Given that we used normalization, this must be compensated by a few sequences with very high numbers of reads (not visible but each counts a lot in the normalization). This suggests that under stress a few regions have many more reads than in controls and as a counterpart, many regions with relatively low number of reads have slightly less reads. By plotting transformed ATAC-Seq read counts of ‘stressed’ vs ‘control’ we see that there are two clouds of points: (i) those regions that have many more reads in stressed than in controls (+1-2 log10units = 10 - 100fold change) and (ii) under the 1:1 stressed-control line those regions that are slightly less represented in the ‘stressed’ than in the ‘controls’ (Figure 6). Regions identified by ChromstaR are present in both clouds (Figure 7).

**Figure 7:**
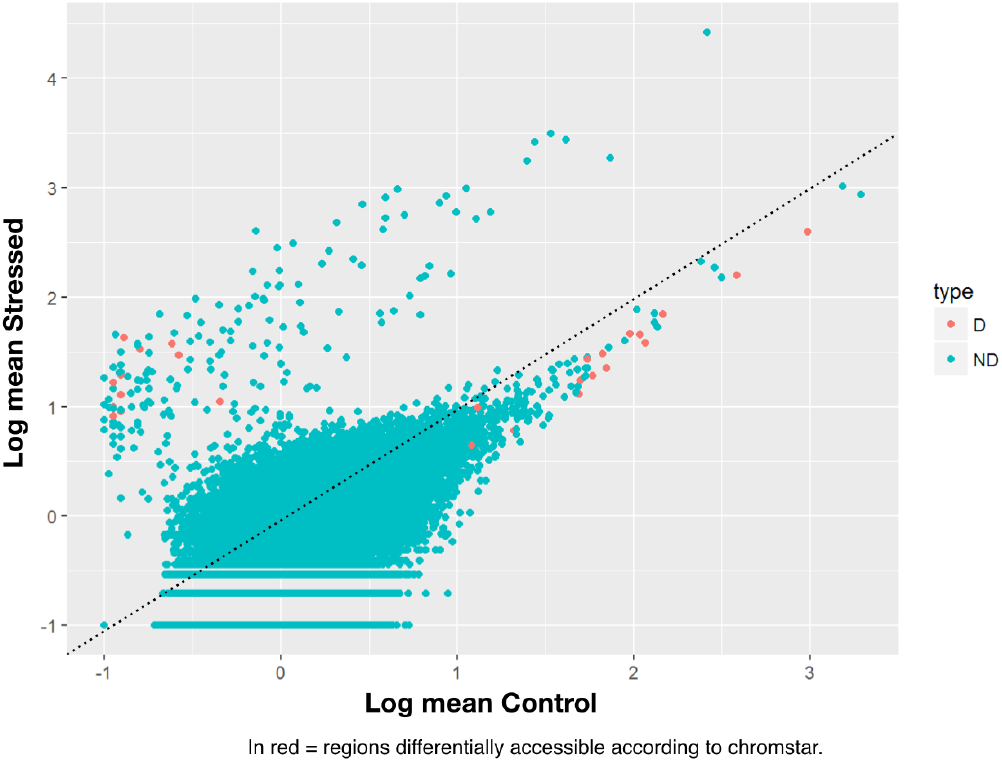
Plot of transformed ATAC-Seq read counts of ‘stressed’ vs ‘control’. Two point clouds of points: 1) at the top-left, those regions have many more reads in stressed than in controls (+1-2 log10units = 10 - 100fold change), and 2) under the 1:1 stressed-control dotted line, those regions that are slightly less represented in the ‘stressed’ than in the ‘controls’. Red dots (D) represent regions that were determined as differently accessible by ChromstaR HMM approach. ND all others.

We therefore conclude that stress modifies the distribution of the numbers of reads drastically: a few hundred regions are much more represented (i.e. Tn5 accessible) in stressed than in control chromatin. This could reflect the fact that under stress these regions become accessible in many more tissues than in control conditions, thus they are captured many more times by the ATAC-seq. As a counterpart, the proportional representation of most other regions slightly decreases (fold change approximately ½ = −0.3 log10 units) but remain generally proportional to their value in controls. Competition for sequencing (PCR amplification selects for the many reads from highly accessible regions) and normalization (divide by the total number of sequences) can be responsible for this.

## Discussion

We report here a very fast and straightforward method to map the chromatin status of individuals using small amounts of input biological material (Augusto *et al.* 2019). The technique is very robust and we have been using it for more than a year now on different species e.g. adult worms of the parasite *Schistosoma mansoni*. The technique was successfully used in the framework of a summer school for field ecologists, some of them with no training in molecular biology. The technique avoids many caveats that are involved with the use of antibody-based methods (Egelhofer *et al.* 2011) and is roughly 6 times faster. In our hands, there was no problem with mitochondrial contamination which is sometimes observed with other ATAC-seq methods. However, it also has some drawbacks: when using aquatic organisms, we observed DNA pollution from other species than the model species/species of interest. It is thus necessary to carefully wash the samples in DNA-free water. In addition, organisms should not be fed a couple of days before the ATAC experiment is performed. Another, inherent weakness of the method is that it provides just a positive readout of Tn5 accessible. These regions and considered nuleosome-free and presumably euchromatic. Absence of ATAC-Seq reads is generally considered as signal for inaccessible, and therefore heterochromatic regions. However, as with any method, the absence of proof is not proof of absence. This caveat is of course shared with any technique that relies on enzymatic accessibility such as DNA footprint, DNAse-seq or FAIRE-seq. It would be desirable to develop techniques that also provide a positive display of heterochromatic regions and without any use of antibodies. Notwithstanding these caveats, due to its minimal training requirements, low starting material as input and price advantage over other techniques, ATAC-seq can be used to develop fast epigenotyping approaches in populations similar to what is done routinely today in population genetics.

We estimate here the epimutation rate to be at least in the order of magnitude of 10^−3^ which is in line to earlier findings (van der Graaf *et al.* 2015) (Roquis *et al.* 2016). However, we also realize that the current analysis methods are not suitable if strong, global changes in chromatin structure occurs. As for genetic analyses, the underlying assumptions of algorithms is that (epi)mutations are rare events. This prompts a need to develop new analysis methods that take large genome wide modifications of the chromatin structure into account. HMM based methods or our transformation method are promising starting points for this purpose.

The development of defensive crests upon exposure to kairomones from predators has been studied in great detail in several *Daphnia* species. It’s developmental dynamics (e.g. (Weiss *et al.* 2015)) and even the genes that are differently expressed (e.g. (Rozenberg *et al.* 2015)) are now known. It is also recognized that genetic, non-genetic and environmental elements interact to bring about phenotypic variation in *daphnia* populations (Harney *et al.* 2017), a phenomenon that is probably applicable to all eukaryotes and that we have recently conceptualized as a systems biology view on inheritance (Cosseau *et al.* 2016). *Daphnia* possess bearers of epigenetic information such as modified histones (Robichaud *et al.* 2012), and DNA methylation (Kvist *et al.* 2018) of the classical mosaic type (Aliaga *et al.* 2019). Despite the the fact that the idea of a *matrix* was known when the term ‘phenotypic plasticity’ was formalised, it remains surprising that, to date, the link between Woltereck’s matrix and the chromatin has not been explicitly made. Maybe because it was too evident, or maybe because the idea that heritable units are composed of several elements and not only DNA fragments (in Woltereck’s words “… *chromosomes are matrix plus gene…*”) is still not entirely accepted by the scientific community. The matrix concept provides a clear key to understanding how organisms can interpret environmental cues and change their phenotype over time spans that are beyond the duration of the cue. In terms of systems biology, the capacity to interpret these cues is the ‘emerging property’ of the inheritance system that is composed of genotype and epigenotype (and potentially cytoplasmic elements and microorganisms). It also allowed eukaryotic cells to generate complexity and thus the symbiontic acquisition of histone-based chromatin organization was probably critical for the evolution of eukaryotic complex cells (Brunk & Martin 2019).

In conclusion, we show here that moderate changes in the environment (during the 20 days, *i.e.* 2-5 generations, from ‘start’ where *daphnia* where introduced into their new water tanks to ‘control’) are accompanied by roughly 4% of epigenetic modifications. When strong environmental cues, such as predator presence, are applied, the chromatin structure is reorganised much more profoundly and many regions become inaccessible to Tn5.

## Materials and Methods

### Daphnia culture and experimental design

A batch of ~300 commercial *Daphnia pulex* was obtained f rom a commercial supplier (Aqualiment: http://www.aqualiment.eu/). At their arrival, daphnia were immediately split into two sets of equal size (~ 150 × 2) and placed in two independent experimental tanks (i.e. initial density of 75 ind.L^−1^), hereafter called the ‘stress’ and the ‘control’ tanks. Each experimental tank consisted in a2-L plastic aquaria (L × l × h = 18 ×12 × 11 cm) supplied with clean water, inside of which a floating plastic fish breeding isolation box (L × l × h = 12,5 × 8 ×7 cm) was placed (Figure 1). These isolation boxes are transparent with a series of 1 mm cracks on the bottom wall to allow water connection between the tanks and inside the isolation box. Daphnia were acclimated in their respective experimental tanks out of the isolation box for 20 days prior to starting the experiment. This lag time before the experiment also allowed the production of new daphnia offspring born in our experimental setup. During this acclimating period, only negligible mortality was observed and newly hatched daphnia were observed in the two experimental tanks. After this 20-day acclimating period, a predator (i.e. a guppy fish previously trained to eat daphnia) was introduced into the isolation box of one experimental tank during 15 days (i.e. hereafter called the ‘stress treatment’, compared to the ‘control treatment’). During the experiment the fish was fed every other day with 10 *daphnia* collected alternatively from the stress and the control tank (i) to avoid subsequent biases in density between the experimental treatments and (ii) to account for a possible effect of daphnia sampling on congeners’ responses. *Daphnia* sampling for fish feeding was achieved using a sterile 3-ml plastic transfer pipet. This experimental setup allowed the daphnia of the stress treatment to experience an indirect predation pressure (i.e. without predation risk) through a direct visual contact with the predator and an olfactory contact with environmental cues released by the predator. Overall the experiment, the *daphnia* and the predator were maintained at room temperature following the natural photoperiod and the former were fed *ad libitum* with clean phytoplancton (i.e. *chlorellasp.*) reared in our lab facilities.

### Sampling and morphometry

Four *daphnia* were sampled during the 20 days acclimating period (called herein ‘start’ population) and immediately processed for ATAC-Seq. At day 15 of stress treatment (2-5 generations), 12 and 14 living *daphnia* were respectively sampled from each of the control and stress treatment by pipetting through a 1 mL automatic pipette with enlarged openings of the pipetting tips and disposed on microscopic slides for dark field microscopy. To avoid experimenter bias 10 different persons sampled at least one control and one stressed *daphnia*. Each *daphnia* was observed and photographed under a stereo microscope (Leica EZ4) at a 100fold magnification using the Leica application suite LAS EZ Version 3.4.0.

From each picture two body lengths were measured (Figure 8): the short length (SL=from the middle of the eye to the base of the apical spine) and the long length (LL=from the middle of the eye to the tip of the apical spine). Finally, each measured animal was then individually transferred to a 1.5 mL Eppendorf tube and was immediately processed for ATAC-seq library preparation. To check for morphological response of *daphnia* to predation pressure we compared the individual ratio of (LL-SL)/SL of each treatment with a student t-test using Excel and http://www.estimationstats.com/#/analyze/two-independent-groups.

**Figure 8:**
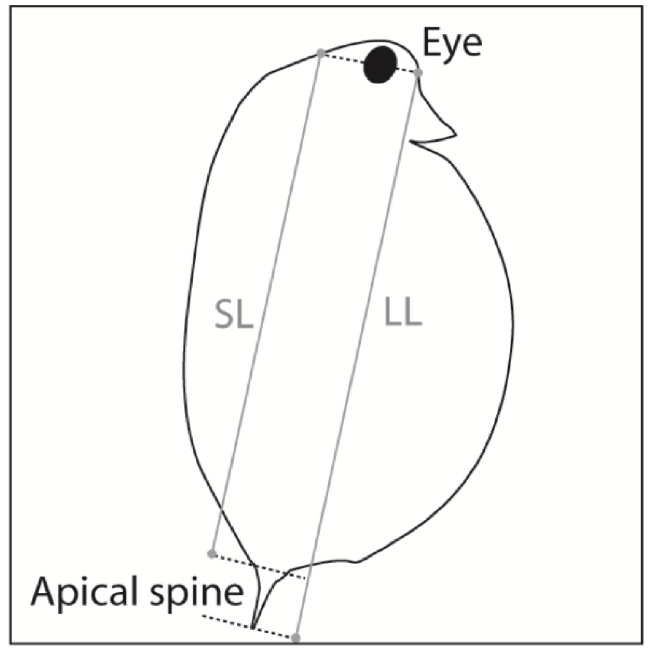
Schematic representation of the measures taken on daphnia. SL = short length, LL = Long length.

### ATAC-Seq

The ATAC-Seq protocol we used is based on Corces et al., (Corces *et al.* 2016) with some modifications (Augusto *et al.* 2019). All water was removed from the d*aphnia* containing tubes through pipetting. *Daphnia* were washed once with 50 μl cold PBS and all liquid was then removed by pipetting. 22 μl nuclease free water, 25 μl 2x TD buffer (Illumina FC-121-1030), 2.5 μl TDE (Tn5 Illumina FC-121-1030) and 0.5 μl 1% IGEPAL CA-630 (Sigma-Aldrich, cat. no. I8896) were added and mixed by pipetting 10 times to disrupt cells. Reaction mixtures were incubated at 37°C for 30 min at 300 rpm agitation. Transposed DNA was immediately purified using a QIAquick PCR Purification Kit (#28106), and purified DNA was eluted into 10 μl of elution buffer (10 mMTris-HCl, pH 8). Libraries were PCR amplified using Promega GoTaq2, universal Ad1_noMX primer and index primer Ad2.* (http://www.nature.com/nmeth/journal/v13/n11/extref/nmeth.3999-S5.xlsx) (each 1.25 μM) that was different for each individual *daphnia*, 10 μl of DNA in a total volume of 50 μl. Pre-amplification was done at 98°C for 30 sec, then five cycles of 98°C for 10 sec, 63°C for 30 sec, 72°C for 1 min. 5 μl of this PCR mixture was used for qPCR analysis to determine the number of additional amplification cycles. Relative fluorescence was plotted versus cycle number and the cycle number that corresponds to one-third of the maximum fluorescent intensity was used for additional PCR amplification. After PCR, size-selection at 300 bp was done on an IP-Star system with Ampure XP beads. Quality and quantity of libraries were checked with an Agilent Bioanalyzer High Sensitivity DNA Assay and library were sequenced on a NextSeq550 High Output Flowcell as paired-end and 75 bp. A detailed step-by-step protocol in Augusto *et al* 2019.

### Detection of chromatin structure differences

Sequence quality was checked with FastQC (http://www.bioinformatics.babraham.ac.uk/projects/fastqc/). Reference genome was downloaded f rom ftp://ftp.ensemblgenomes.org/pub/metazoa/release-40/fasta/daphnia_pulex/dna/Daphnia_pulex.V1.0.dna.toplevel.fa.gz, corresponding to Gen Bank assembly accession GCA_000187875.1. Alignment was done with Bowtie2 evoking the following parameters: bowtie2-align-s basic-0 -p 6 -x genome -N 1 -L 32 -i S,1,1.15 --n-ceil L,0,0.15 --dpad 15 --gbar 4 --end-to-end --score-min L,−0.6,−0.6. Uniquely aligned reads were retained by filtering the tag “XS:i:” that is absent in their alignement annotations.

For visualisation of ATAC profiles all BAM files for each condition were merged, converted to header-free SAM, and downsampled with a custom script that draws random lines to 409,000 aligned reads. This corresponds to the condition with the lowest number of aligned reads. For analysis of individual *daphnia* PCR duplicates were removed with SamTools RmDup. Bedgraph files were generated with MACS2 using model building, lower fold bound of 5, upper fold bound 50, band width 300 bp, minimum FDR for peak detection of 0.05, an effective genome size of 150,000,000, and without calling broad regions. Bedgraphs were loaded into IGV for visual inspection. For analysis of individual *daphnia*, background correction was done with MACS bdgcmp. Bedgraph was converted into BigWig. The DeepTools suite was used for representation of metagene profiles based on this over 15,287 genes on the forward strand. Gene annotation files were downloaded from ftp://ftp.ensemblgenomes.org/pub/metazoa/release-40/fasta/daphnia_pulex/cds/Daphnia_pulex.V1.0.cds.all.fa.gz. More information is available at https://metazoa.ensembl.org/Daphnia_pulex/Info/Annotation/

Two different approaches were used for further data analysis. One uses a combination of peakcalling with MACS2, extraction of read coverage in peaks with BEDtools, and DESeq2 for differential analysis. To detect all peak regions for all conditions, BAM files of control and stress conditions were merged and peakcalling was performed with MACS2 as described above. The number of reads overlapping peak regions was extracted with bedtools intersect -a peakfile.bed -b individual_bam_files.bam-header -wa -c, Columns 4 and 11, corresponding to peak-names and number of overlapping features, i.e. coverage were used as input for DESeq2. All analyses were done at the galaxy instance of the Labex CeMEB/IHPE (http://bioinfo.univ-perp.fr).

The second approach was based on Hidden-Markow-Models (HMM) implemented in ChromstaR (v.1.2.0) for genome-wide characterization of open chromatin landscape. On this approach control and stress condition were processed in two steps: (1) we fitted a univariate HMM over each ATAC-seq samples individually and (2) we performed a multivariate HMM over the combined ATAC-seq samples in each condition. For that, BAM files were processed under the differential mode, with a false discovery rate (FDR) cutoff of 0.05 and bin size of 500.

## End Matter

### Author Contributions and Notes

R.A. and C.G. designed research, all authors performed research, P.D. wrote software, R.A., P.D. and C.G. analyzed data; and all authors wrote the paper.

The authors declare no conflict of interest.

This article contains supporting information online on NCBI SRA.

## Acknowledgments

These experiments were done in the framework of the RTP 3E summer school “Epigenetics for field ecologists” and received support from the CNRS.

