## Supplementary figure 1 for "Chromatin structure changes in *Daphnia* populations upon exposure to environmental cues – or – The discovery of Wolterecks “Matrix”"

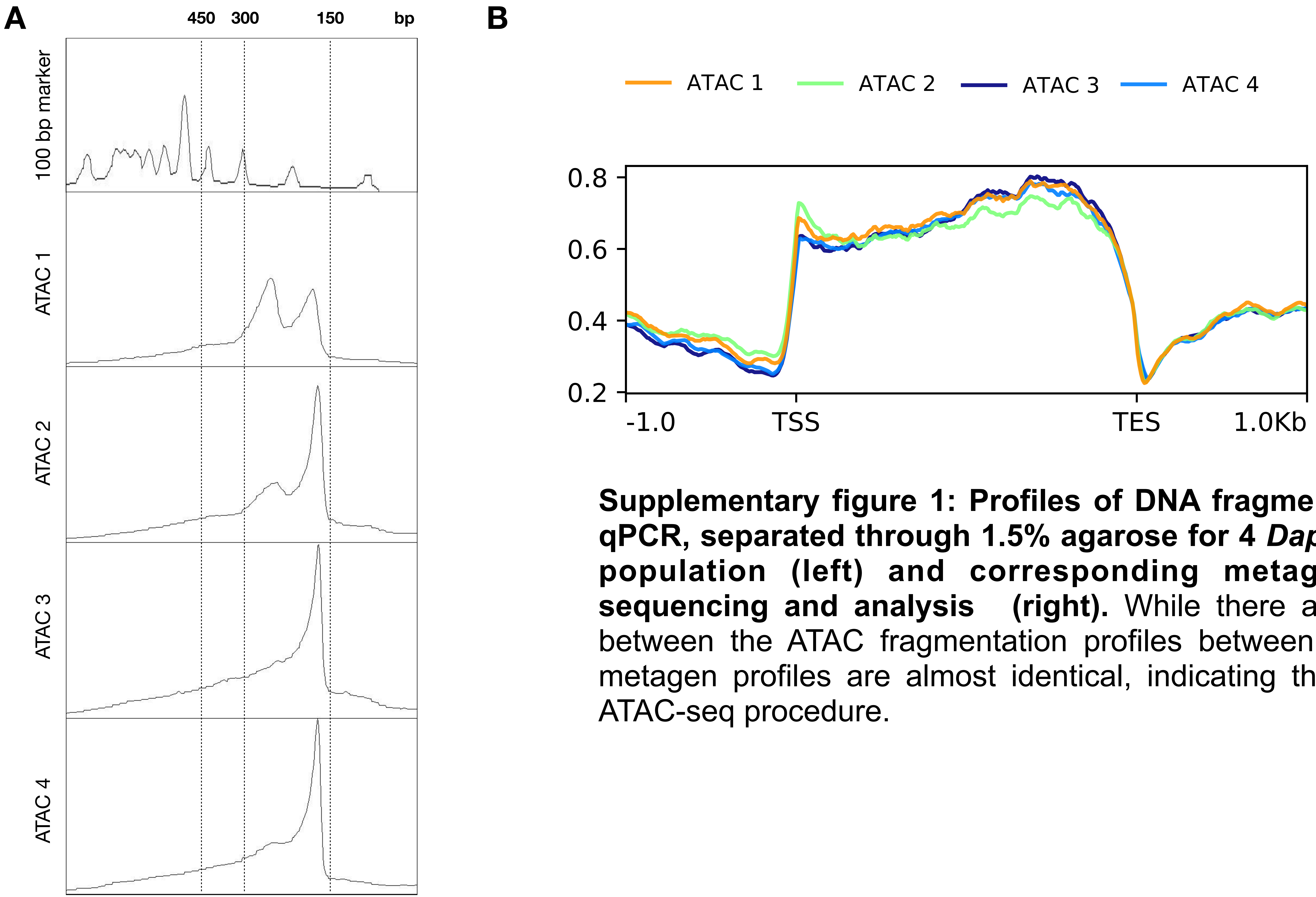


**Supplementary figure 1: Profiles of DNA fragments after ATAC and qPCR, separated through 1.5% agarose for 4 *Daphnia* from the start population (left) and corresponding metagen profiles after sequencing and analysis (right).** While there are small differences between the ATAC fragmentation profiles between individuals (A), the metagen profiles are almost identical, indicating the robustness of the ATAC-seq procedure.
