## Supplementary figure 2 for "Chromatin structure changes in *Daphnia* populations upon exposure to environmental cues – or – The discovery of Wolterecks “Matrix”"

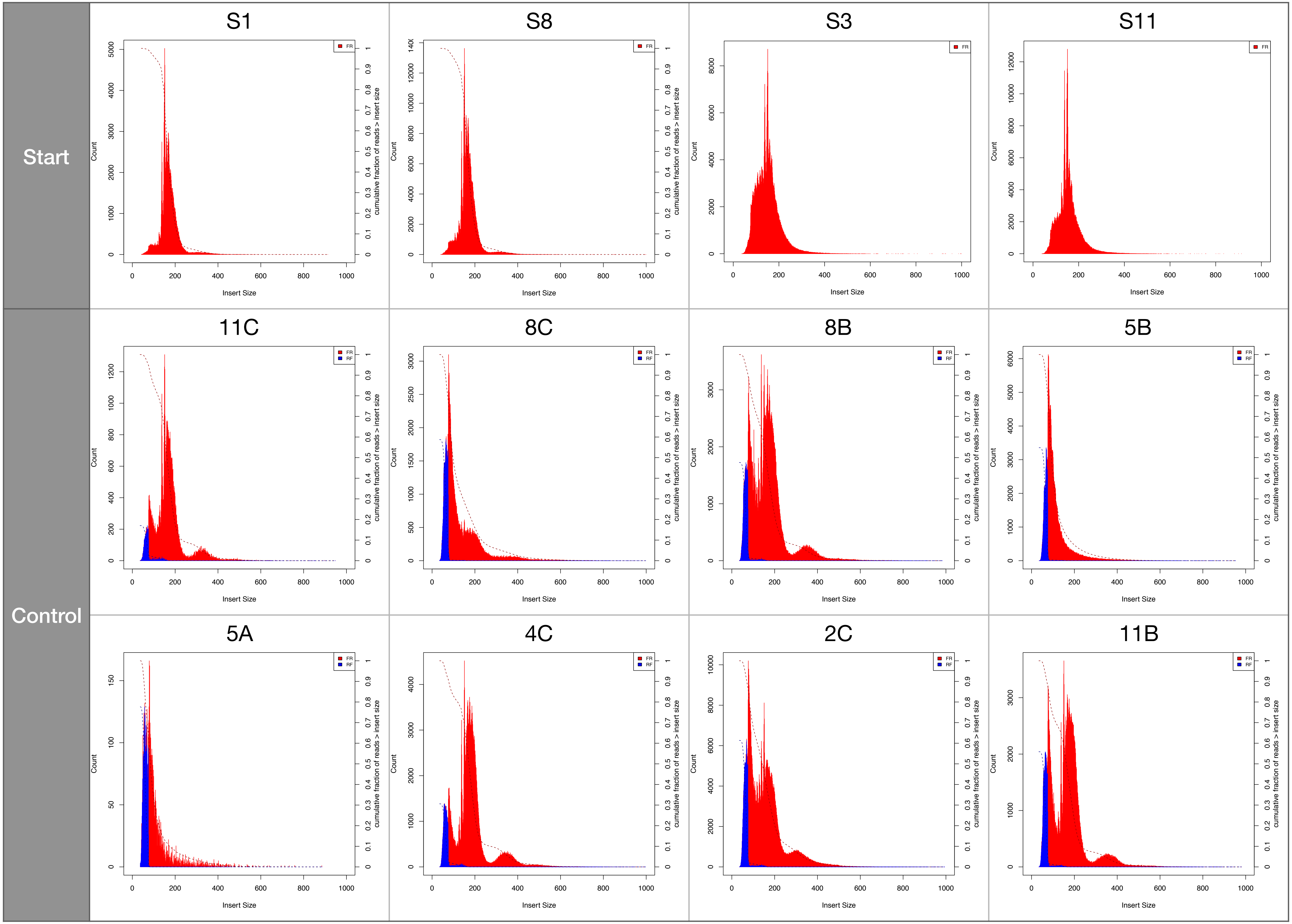

Start
Control
S1
S8
S3
S11
11C
8C
8B
5B
5A
4C
2C
11B
FR
RF
Count
Insert Size
cumulative fraction of reads > insert size

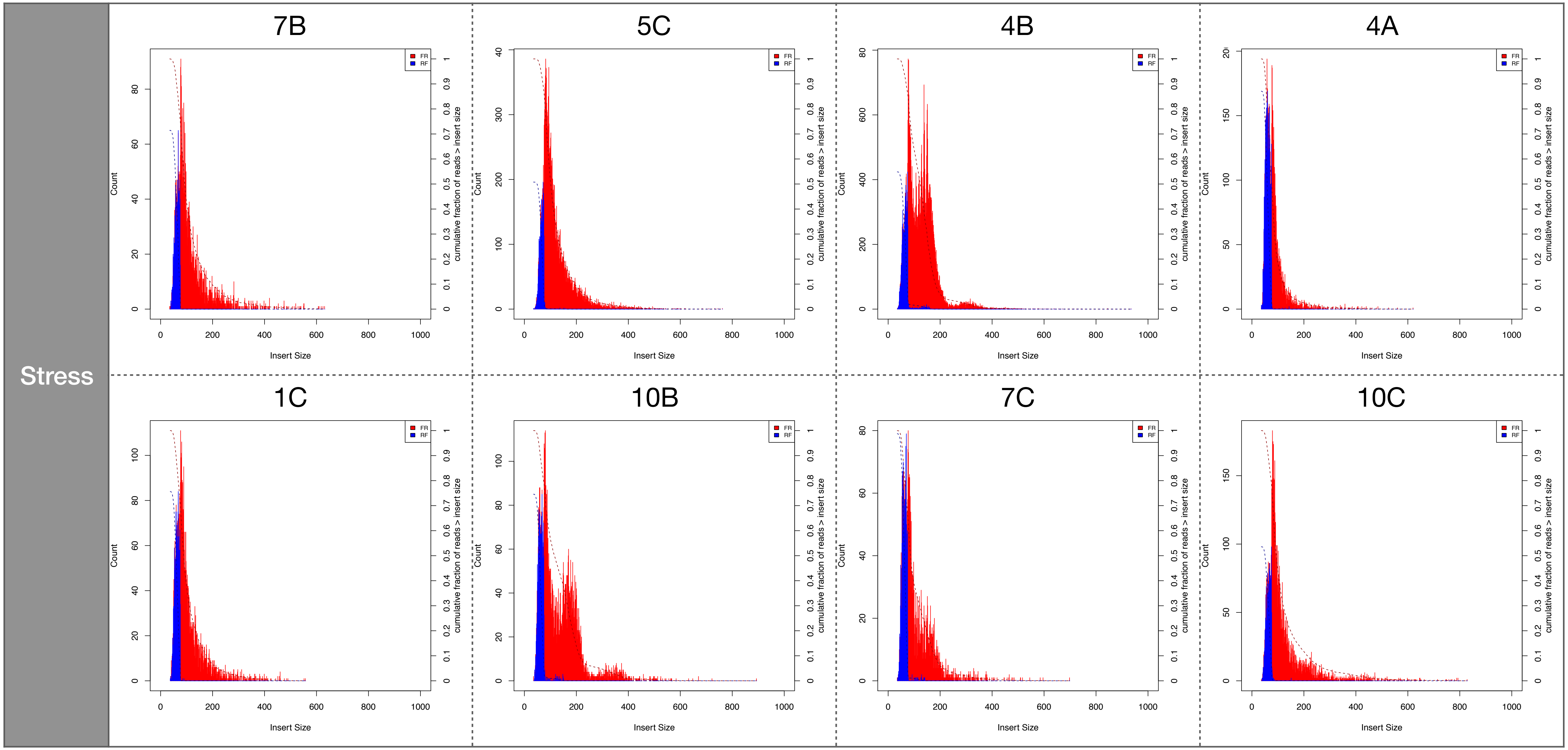


**Supplementary figure 2: Insert size distributions for start, control and stress population.** Every box library for one individual.
